# Pyroptosis leads to loss of centrosomal integrity in macrophages

**DOI:** 10.1101/2023.11.22.568260

**Authors:** Siyi Bai, Fatima Martin-Sanchez, David Brough, Gloria Lopez-Castejon

## Abstract

NLRP3 forms a multiprotein inflammasome complex to initiate the inflammatory response when macrophages sense infection or tissue damage, which leads to caspase-1 activation and maturation and release of the inflammatory cytokines interleukin-1β (IL-1β) and IL-18, and Gasdermin-D (GSDMD) mediated pyroptosis. NLRP3 inflammasome activity must be controlled as unregulated and chronic inflammation underlies inflammatory and autoimmune diseases. Several findings uncovered that NLRP3 inflammasome activity is under the regulation of centrosome localized proteins such as NEK7 and HDAC6, however, whether the centrosome composition or structure is altered during the inflammasome activation is not known. Our data show that levels of the centrosomal scaffold protein pericentrin (PCNT) are reduced upon NLRP3 inflammasome activation via different activators in human and murine macrophages. PCNT loss occurs in the presence of membrane stabilizer punicalagin, suggesting this is not a consequence of membrane rupture. We found that PCNT loss is dependent on NLRP3 and active caspases as MCC950 and pan caspase inhibitor ZVAD prevent its degradation. Moreover, caspase-1 and GSDMD are both required for this NLRP3-mediated PCNT loss because absence of caspase-1 or GSDMD triggers an alternative regulation of PCNT via its cleavage by caspase-3 in response to nigericin stimulation. PCNT degradation occurs in response to nigericin, but also other NLRP3 activators including lysomotropic agent L-Leucyl-L-Leucine methyl ester (LLOMe) and hypotonicity. Our work reveals that the NLRP3 inflammasome activation affects centrosome composition and structure which may deepen our understandings of how activated NLRP3 inflammasomes are involved in the pathogenesis of inflammatory diseases.

## Introduction

Inflammasome activation in macrophages is an early event occurring at the initiation of an inflammatory response. Inflammasomes, formed by sensor proteins such as NLRP3, assemble in response to pathogenic or damage signals leading to the activation of caspase-1 which is responsible for the cleavage of pro-IL1β and pro-IL18 into their active forms [1, 2] as well as for the cleavage of GSDMD, which N-terminal domain forms pores in the membrane required for IL-1β and IL-18 release [3]. GSDMD cleavage is also required for the induction of pyroptosis, a programmed lytic cell death induced upon inflammasome-caspase-1 activation [4].

Subcellular localization of NLRP3 inflammasome is important for its function. Several organelles contribute to NLRP3 regulation and activation either by acting as assembly platforms or as sensors of NLRP3 activating stimuli that mediate inflammasome assembly [5]. The centrosome is one of such organelles. Centrosome associated proteins MARK4, HDAC6, NEK7, PLK1 and PLK4 contribute to either the trafficking of NLRP3 to the centrosome or to regulation of NLRP3, mediating its activation [6,7,8,9,10]. However, little is known about how changes to the centrosome composition and integrity are related to inflammasome activation.

The centrosome is a dynamic organelle which plays important roles in microtubule organisation and in cell cycle but also in stress and damage responses [11, 12]. Centrosomes are formed by the centrioles and a cloud of proteins around them called the pericentriolar material (PCM) [13]. Pericentrin (PCNT) is one of the main components of the PCM of the centrosome. In humans it presents two main isoforms PCNT-A (220 kDa) and PCNT-B (also known as kendrin, 340 kDa) and that shares its N-terminal end with PCNT-A [14]. The composition of the centrosome is altered during the cell cycle [15] as well as in response to stresses such as DNA damage [16] or heat shock [17]. Sensing of pro-inflammatory stimuli including the bacterial component LPS induces cell cycle arrest in G1 [18]. Moreover, LPS increases the accumulation of pericentriolar components at the PCM including PCNT and γ-tubulin reflecting an alteration of the PCM [19]. Interestingly monocytes from febrile patients with a fever or subjected to heat stress present a reversible loss of integrity of the centrosome [17].

Pyroptosis and apoptosis are different forms of programmed cell death. Pyroptosis, unlike apoptosis, does not depend on apoptotic caspases (caspase-3, -9, etc.) activation [20] but on proinflammatory caspases (e.g., caspase-1) and the assembly of the GSDMD pore at the plasma membrane [21]. Pyroptosis is a lytic cell death where contents are released to the extracellular environment and contribute to inflammation [22]. It is become clear however, that in the absence of GSDMD and hence pyroptosis, caspase-1 activation triggered by NLRP3 inflammasome leads to caspase-3 activation and consequently apoptosis [23, 24]. This process is followed by caspase-3 mediated cleavage of Gasdermin E, that eventually also leads to a pyroptotic event [23, 24]. Moreover, cells lacking caspase-1 are also able to trigger incomplete pyroptosis via caspase-3, highlighting a tight cross-regulation between these two types of cell death and caspases [25].

The centrosome can assemble and disassemble in response to specific stimuli and this is essential for appropriate cell division. Here, PCNT is cleaved by separase, a member of the Cell Death family of cysteine proteases, which also includes caspases [26]. PCNT cleavage by separase occurs at R2231 (leading to a 275kDa fragment), and mutations in this site suppresses centriole disengagement and subsequent centriole duplication [27]. Cleavage of PCNT can also be mediated by caspase-3 in response to apoptotic stimuli. Seo and Rhee (2018) demonstrated that treatment of HeLa cells with apoptotic agents led to the cleavage of PCNT while no cleavage of γ-tubulin was detected [28]. The specific PCNT cleavage sites targeted by caspase-3 remain to be identified. However, it remains unclear whether the centrosome is targeted by caspase-1 in pyroptotic cells and how this is governed.

Here, we investigated the relationship between centrosome and inflammasome activation in macrophages. We found that centrosome structure and PCNT protein are lost in response to NLRP3 activators nigericin, LLOMe, and hyptonocity and that this is dependent on the NLRP3/caspase-1/GSDMD signalling axis. This work highlights that the centrosome is altered and dysfunctional in the pyroptotic environment formed by NLRP3 inflammasome activation.

## Materials and Methods

### Reagents and antibodies

LPS (*Escherichia coli* 026:B6); Nigericin (N7143); protease inhibitor cocktail (P8340); phorbol 12-myristate 13-acetate (PMA, P8139); penicillin–streptomycin (Pen/Strep, P4333); Punicalagin (P0023); Caspase-1 Inhibitor IV (YVAD, 400015); MG-132 (M7449); Bovine Serum Albumin (A7906) and Formaldehyde solution (50-00-0) were from Sigma. Dulbecco’s Phosphate Buffered Saline (PBS, D8537); DAPI (28718-90-3); Pepstatin A Methyl Ester (Pepstatin A, 516485) and MCC950 (256373-96-3) were purchased from Merck. Ultrapure™ DNase/RNase-Free Distilled Water (10977035) was from Invitrogen. Zeocin (J67140-8) and MeOH (67-56-1) was sourced from Thermo Scientific. Foetal bovine serum (FBS, S181H-500) was from Gibco. Fluorescence Mounting Medium (S3023) was obtained from Agilent. CA-074 methyl ester (CA-074, S7420) was from Selleckchem. Z-VAD-FMK (ZVAD, 001) was from R&D Systems. Z-DEVD-FMK was obtained from APExBIO. E-64-D (BML-PI107) was sourced from Enzo Life Sciences. L-Leucyl-L-Leucine methyl ester (LLOMe, 16008) was from Cayman. DMSO (7726) was from Bio-Techne.

Primary antibodies used for Western blot assays were as follows: anti-pericentrin (1:500, rabbit polyclonal, Abcam, ab4448), anti-pericentrin (1:200, rabbit polyclonal, Invitrogen, PA5-115736), anti-γ-tubulin (1:1000, mouse monoclonal, Merck, T6557), anti-caspase-1 p20 (1:500, rabbit monoclonal, Cell Signalling Technology, 3866), anti-caspase-3 (1:500, rabbit monoclonal, Abcam, ab32351), anti-NLRP3 (1 μg/ml, mouse monoclonal, Adipogen, AG-20B-0014) and anti-β-actin-HRP (0.2 μg/ml, mouse monoclonal, Sigma, A3854). HRP conjugated secondary antibodies used for Western blot were anti-rabbit-HRP (0.25 μg/ml, goat polyclonal, Dako, P0448) and anti-mouse-HRP (1.3 μg/ml, rabbit polyclonal, Dako, P0260).

Primary antibodies used for immunofluorescence were: anti-pericentrin (rabbit polyclonal, Abcam, ab4448), anti-pericentrin (rabbit polyclonal, Invitrogen, PA5-115736), anti-ASC (mouse, Biolegend, 676502), anti-ASC (rabbit polyclonal, AdipoGen Life Science, AG-25B-0006-C100), anti-ninein (mouse monoclonal, Santa Cruz, sc-376420), Anti-NLRP3 (mouse monoclonal, AdipoGen Life Science, AG-20B-0014) at 1:1000 dilution or anti-γ-tubulin (mouse monoclonal, Merck, T6557) at 1:500 dilution.

### Cell Culture and treatments

THP1^ATCC^, THP1^Caspase1-/-^ and THP1^GSDMD-/-^ cells were maintained in complete RPMI-1640 (with 2 mM L-glutamine, 10% FBS and Pen/Strep (100 U/ml)). THP1^Null2^ and THP1^NLRP3^ ^PYD^ deficient cells (THP1^NLRP3-/-^) were also cultured in complete RPMI-1640 plus Zeocin (100 μg/mL). THP1^ATCC^ cell line was sourced from ATCC. THP1^Null2^ and THP1^NLRP3-/-^ cell lines were purchased from Invivogen. THP1^Caspase1-/-^ cells were a gift from Prof Veit Hornung (Ludwig Maximilian University of Munich). THP1^GFP-NLRP3^ and THP1^GSDMD-/-^ cells were generated in the Lopez-Castejon’s lab as previously described [29, 35]. All cultures were maintained in humidified incubators at 37 °C, 5% CO_2_.

For Bone Marrow Derived macrophages (BMDMs) isolation, femur and tibia from 6-8 months old C57BL/6J mice were removed. Bone marrow was flushed out, resuspended in DMEM supplemented with 20% L929 supernatant, 10% FBS and 1% Pen/Strep (100 U/ml), then cultured for 6 days until differentiation into macrophages. The resulting BMDMs were detached with cold PBS and seeded on cell culture plates for use next day.

THP1 cells were plated at a density of 1×10^6^ cells/ml and differentiated with 0.05 μM PMA. After 24 h, media were removed and replaced with fresh media. Experiments were carried out the following day. During stimulation, cells were kept in E-total buffer (147 mM NaCl, 10 mM Hepes, 10 mM D-glucose, 2 mM KCl, 2 mM CaCl_2_, 1 mM MgCl_2_, buffered to pH 7.4).

### Cell death Assay

Cell death was measured using quantitative assessment for the release of lactate dehydrogenase (LDH) into cell supernatants, after a centrifugation step of 1 min at 13,000×g at 4°C, to remove any dead/floating cells. CytoTox 96® Non-Radioactive Cytotoxicity Assay (Promega, G1780) was used according to the manufacturer’s instructions. Absorbance values were recorded at 490 nm and the results were expressed as a percentage of LDH release relative to the total cells lysed.

### Caspase-1 activity Assay

Caspase-1 activity was measured in the supernatants using Caspase-Glo® 1 Inflammasome Assay (Promega G9951). Briefly, cell supernatants were combined with Z-WEHD aminoluciferin substrate for 0.5 h before recording luminescence. The results were expressed as a fold change relative to untreated cells.

### Cathepsins activity Assay

The activity of cathepsin B and cathepsin D was measured using Abcam Fluorometric Activity Assay Kits (ab65300 and ab65302 for cathepsin B and cathepsin D, respectively). Briefly, cell lysates were incubated with reaction mix including reaction substrate and buffer at 37 °C for 90 min following the manufacturer’s instructions. Fold change from the untreated cells control was calculated for all experimental groups.

### Proteasome activity Assay

Proteasome activity was measured in cell lysates using the Proteasome-Glo™ 3 Substrate System (Promega, G8531). Corresponding reagents for testing as chymotrypsin-like, trypsin-like, and caspase-like activity of the of proteasome are included in this kit. Manufacturer’s instructions were followed. 30 min after adding the individual Proteasome-Glo™ Reagents separately, luminescence was recorded as relative light units (RLU) on a GloMax® 96 Microplate Luminometer.

### Enzyme-Linked Immunosorbent Assay (ELISA)

Levels of human IL-18 and mouse IL-1β were measured in the cell supernatants using ELISA kits from R&D Systems (DY318) and (DY401-05), respectively. ELISAs were performed following the manufacturer’s instructions.

### Western Blot

Cells were lysed for at least 20 min on ice using a RIPA lysis buffer (50 mM Tris–HCl, pH 8, 150 mM NaCl, 1% NP-40, 0.5% sodium deoxycholate and 0.1% sodium dodecyl sulphate, SDS), supplemented with a protease inhibitor cocktail (1:100). Lysates were then centrifuged at 13,000 ×g for 10 min to remove the insoluble fraction. Protein concentration was measured by BCA assays (Thermo Scientific Pierce, 23225), following the manufacturer’s guidelines, and an equal amount of protein was loaded for each sample. Cell supernatants were centrifuged at 500 ×g for 5 min to remove dead cells and concentrated with 10 kDa MW cut-off filters (Amicon, Merck Millipore), as described by the manufacturer. In cases where the whole well lysate was assayed, the cells were directly lysed in the well by the addition of 1% (vol/vol) Triton X100 with a protease inhibitor cocktail (1:100). Whole well lysates were then centrifuged at 21,000 xg for 10 min to remove the insoluble fraction. Lysates, supernatants and whole well lysates were diluted in Laemmli buffer containing 1% 2-mercaptoethanol, heated at 95°C for 10 min and resolved by SDS–PAGE.

Separated proteins were transferred onto nitrocellulose membranes and blocked in 5% Milk PBS-Tween (0.1%) for 1 h at room temperature (RT). Membranes were then incubated with the specific primary antibody in blocking buffer for 1 h at RT. Then, membranes were washed three times in PBS-Tween (PBS-T, 0.1%) for 10 min per wash and subsequently incubated for 1 h at RT with a horseradish peroxidase-conjugated secondary antibody. Membranes were then washed as before and visualised using Clarity™ Western ECL Blotting Substrate (Bio-Rad, 1705061) in a ChemiDoc™ MP Imager (Bio-Rad). Semiquantitative densitometry analysis of the western blot for PCNT were performed using ImageJ.

### Immunofluorescence

PMA-differentiated THP1 seeded and stimulated on coverslips were fixed with 4% formaldehyde for 10 min at RT, and followed by ice-cold 100% MeOH permeabilization at -20°C for 10 min. Cells were blocked for 30 min with 1% BSA previous 1h incubation at RT with the primary antibody rabbit anti-pericentrin (Abcam, ab4448), rabbit anti-pericentrin (Invitrogen, PA5-115736), mouse anti-ASC (Biolegend, 676502), rabbit anti-ASC (AdipoGen Life Science, AG-25B-0006-C100), mouse anti-ninein (Santa Cruz, sc-376420) and mouse anti NLRP3 (AdipoGen Life Science, AG-20B-0014) at 1:1000 dilution or mouse anti-γ-tubulin (Merck, T6557) at 1:500 dilution. Coverslips were then incubated for 1h at RT with the appropriate Alexa Fluor conjugated secondary antibody (Invitrogen, 1:300 dilution), incubated for 10 min with DAPI at 1μg/mL in PBS and mounted on slides using Dako Fluorescence Mounting Media.

For quantification, images were acquired on an Olympus IX83 inverted microscope using UV (395 mM), Cyan (470 nm) and red (640 nm) Lumencor LED excitation, a 20x UPlanSApo (oil) objective and the Sedat QUAD filter set (Chroma [89000]). The images were collected using a R6 (Qimaging) CCD camera with a Z optical spacing of [0.2 μm]. Maximum intensity projections are shown in the results. Four different fields per image (200-300 total cells per condition) were used in quantification. Number of cells positive for the protein of interest (in ImageJ) was counted and expressed relative to total number of nuclei as an indication of total number of cells.

### Statistical Analysis

GraphPad Prism 9 software was used to carry out all statistical analysis. One-way ANOVA with the Dunnett’s test or two-way ANOVA with the Tukey’s test were applied in multiple comparisons. Data are shown as mean +/- standard deviation (S.D.). ns is considered as not statistically significant. *p < 0.05, **p < 0.01, ***p < 0.001, ****p < 0.001.

## Results

### Nigericin treatment of THP-1 cells results in loss of centrosomal integrity

To investigate whether NLRP3 activation influences the integrity of the centrosome we used PMA-differentiated THP1 cells and treated them with the well-known NLRP3 activator nigericin [30] at time points as indicated. We have previously shown that LPS priming has minimal effect on NLRP3 inflammasome activation in THP1 cells and to reduce complexity in the experimental system, we did not LPS-prime THP1 cells here. We found that nigericin induced cell death, caspase-1 activation as well as IL-18 release overtime (Fig. 1A-C). We then looked at the expression of PCNT in cell lysates by Western blot. We detected three main different bands for PCNT; PCNT-A (220 kDa) and PCNT-B (340 kDa) as well as a band corresponding to the separase-cleaved PCNT-B (275kDa) in untreated cells. We found that levels of PCNT started decreasing after 15 min of nigericin treatment (Fig. 1D). Expression of another PCM component, γ-tubulin was however unchanged as was β-actin which was used as loading control (Fig. 1D). To determine if the decrease in PCNT levels were due to its release we concentrated the supernatants from those cells and ran western blots for the same three proteins. We could not detect PCNT in these supernatants however γ-tubulin and β-actin were present after 30 min of treatment corresponding to an increase in cell death and protein release (Fig. 1E). To further examine that this reduction in PCNT levels was not due to the release of this protein we performed the same nigericin time-response experiment but collected lysates and supernatants together (whole well lysate). Here we found again that PCNT levels decreased over time after nigericin treatment while γ-tubulin and β-actin levels remained unchanged (Fig. 1F).

**Fig. 1.**
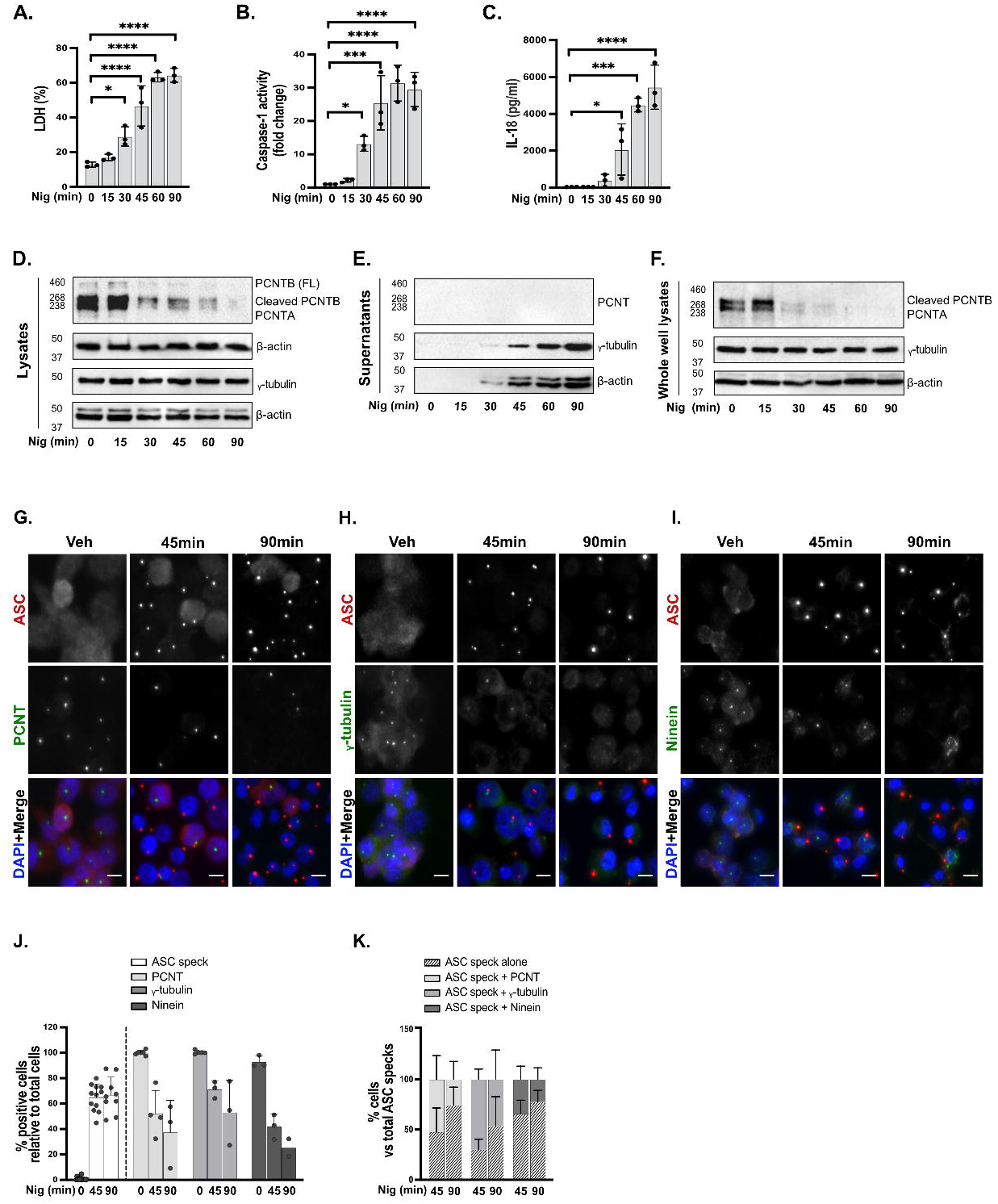
Nigericin treatment of THP-1 cells results in loss of centrosomal integrity. THP1^ATCC^ cells were left untreated or treated with nigericin (10 µM) at the indicated time points to activate the NLRP3 inflammasome, then cell lysates and supernatants were analyzed for PCNT, γ-tubulin, and β-actin expression by western blot (A-F, N=3). Lysates were analyzed for PCNT and γ-tubulin as well as loading control β-actin (42 kDa) (D), and supernatants (E) and whole well lysates (F) were also analyzed for those proteins. Cell death was measured by LDH assay and shown as percentage relative to total cell death (A). Caspase-1 activity in supernatants was measured and shown as fold change relative to control (B). IL-18 in the supernatants was measured by ELISA (C). THP1^ATCC^ cells were stimulated with nigericin (10 µM) for 45 min or 90 min. (G-K, N=3). Immunofluorescence was used to analyze the centrosomal proteins including PCNT (G), γ-tubulin (H) and ninein (I) as well as ASC to determine the NLRP3 inflammasome activation. Percentages of ASC speck, PCNT, γ-tubulin or ninein positive cells relative to total cells (J) and PCNT, γ-tubulin or ninein positive cells in ASC positive cells (K) were quantified by the Image J. Independent experiments. For multiple comparisons, one-way ANOVA with the Dunnett’s test for time response in THP1^ATCC^ cells was applied. Data was shown as mean ± S.D., *p < 0.05, **p < 0.01, ***p < 0.001, ****p < 0.001 were considered statistically significant.

To further investigate the effect of nigericin on centrosomal integrity we performed immunofluorescence on PMA differentiated THP1 cells, in response to nigericin treatment for 45 and 90 min, for the PCM proteins PCNT, γ-tubulin, and the centriole distal appendix protein ninein [31] and ASC, to determine inflammasome activation (Fig. 1G-K). In line with our western blot data, we found that PCNT centrosomal signal was lost after nigericin treatment (Fig. 1G, J). A decrease in centrosomal γ-tubulin and ninein stain was also observed (Fig. 1H, I, J). We found that centrosomal loss was mainly observed in cells that presented ASC specks, indicating that the ASC-speck remains intact in cells despite loss of centrosome integrity (Fig. 1K). This indicates that nigericin treatment and inflammasome activation leads to loss of centrosomal integrity in macrophages.

### Centrosomal disorganization triggered by nigericin is NLRP3 dependent

To determine if this PCNT loss was dependent on NLRP3 we treated PMA-differentiated THP1 cells either WT (parental THP1^Null2^) or expressing an endogenous PYD-deficient NLRP3 (THP1^NLRP3-/-^) with nigericin (10 µM, 45 min). We found that PCNT protein level was reduced in response to nigericin in the parental THP1 cell line and that this did not occur in THP1^NLRP3-/-^ (Fig. 2A-C). Nigericin treatment of THP1^NLRP3-/-^ cells did not result in increased cell death, caspase1 activation or IL-18 release (Fig. 2D, Fig. S1A, B) after nigericin treatment unlike WT cells, confirming deficient function of NLRP3 inflammasome. These results suggest that centrosome is perturbed after the NLRP3 inflammasome is activated.

**Fig. 2.**
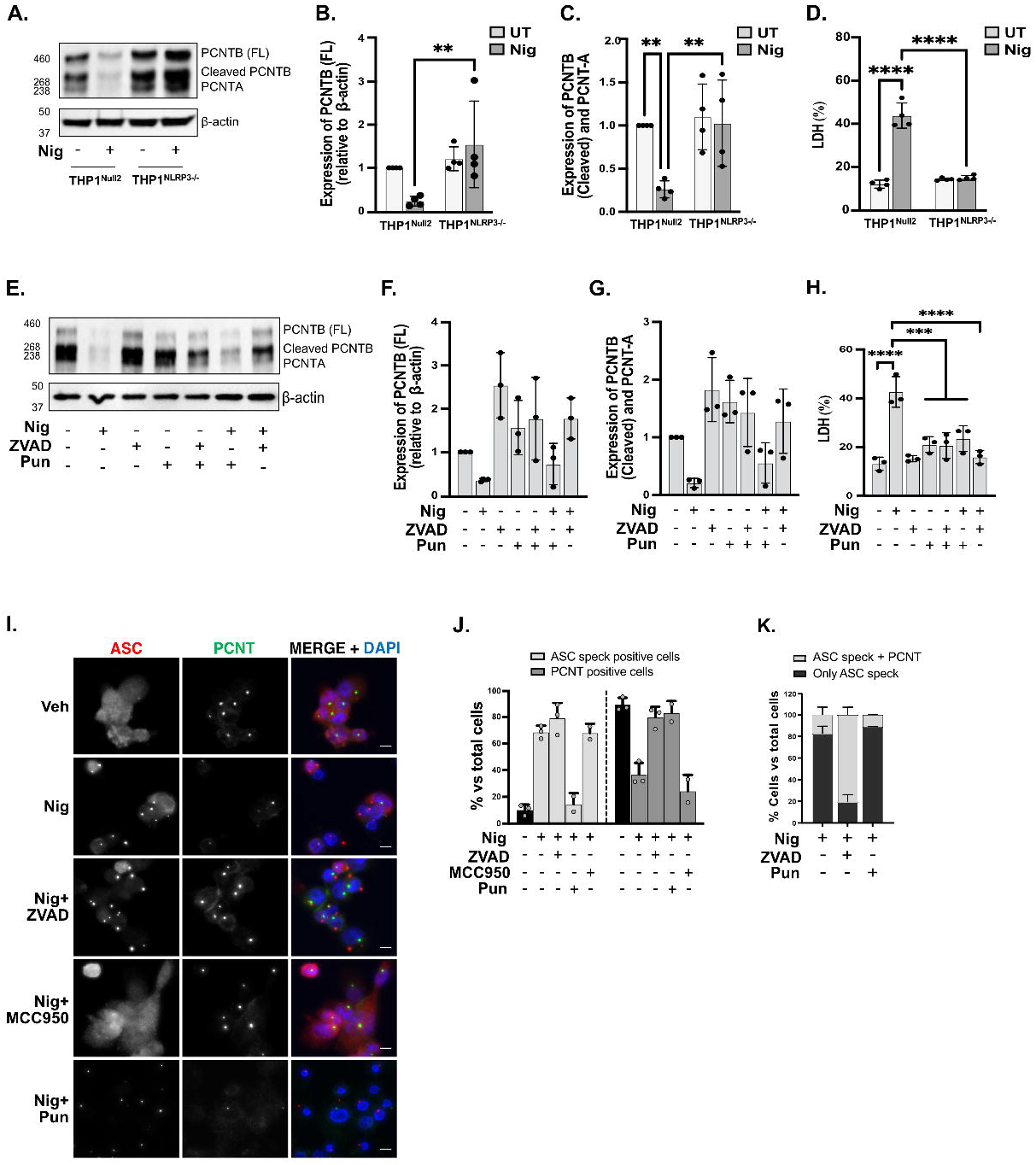
Centrosomal disorganization triggered by nigericin is NLRP3 dependent. THP1^Null2^ and THP1^NLRP3-/-^ cells were stimulated with nigericin (10 µM, 45 min) (A-D, N=4). Lysates were analyzed for PCNT as well as loading control β-actin by western blot (A). Densitometry analysis of relative expression of full length PCNT-B (340 kDa) (B) and cleaved PCNT-B (275kDa)/PCNT-A (220kDa) (C) compared to the control (β-actin) by the Image Lab 6.0. Cell death was measured as described above and shown as percentage relative to total cell death (D). THP1^Null2^ (E-H) or THP1^ATCC^ (I-K) cells were left untreated or treated with punicalagin (50 µM, 15 min) prior to treatment with ZVAD (50 µM, 40 min), after which cells were stimulated with nigericin (10 µM, 45 min) to activate the NLRP3 inflammasome (N=3). Lysates were analyzed for PCNT expression as well as loading control β-actin (E). Relative expression of full length PCNT-B (F) and cleaved PCNT-B/PCNT-A (G) compared to the control were quantified as described above. Cell death was measured and shown (H). Immunofluorescence was used to analyze PCNT and ASC (I). Percentages of PCNT or ASC speck positive cells relative to total cells (J) and both PCNT and ASC positive cells or only ASC speck positive cells in total cells (K) were quantified by the Image J. Independent experiments. For multiple comparisons, one-way ANOVA with the Dunnett’s test in THP1^Null2^ (E-H) and two way ANOVA with the Tukey’s test for comparing nigericin treated THP1^Null2^ and THP1^NLRP3-/-^ were applied. Data was shown as mean ± S.D., *p < 0.05, **p < 0.01, ***p < 0.001, ****p < 0.001 were considered statistically significant.

To further discount that our observation on PCNT downregulation was due to its release due to pyroptosis, we used punicalagin (50 µM, 15 min) to inhibit membrane permeability as punicalagin allows for NLRP3 inflammasome activation but not IL-18 release [29]. We also pre-treated THP1 cells with ZVAD a pan caspase inhibitor to prevent consequences of NLRP3 activation (caspase activation and IL-18 release). We found that, while ZVAD prevented loss of PCNT in response to nigericin, punicalagin treatment did not prevent loss of PCNT in response to NLRP3 activation (Fig. 2E-G). This is despite of punicalagin and ZVAD inducing a reduction in cell death (Fig. 2H) as well as in active caspase-1 and IL-18 release in response to nigericin (Fig. S1C, D). We further confirmed these results by measuring PCNT signal and ASC-speck formation by immunofluorescence after nigericin treatment in the presence of ZVAD, punicalagin and the NLRP3 inhibitor MCC950 (Fig. 2I-K). As before we found that centrosomal loss after nigericin treatment was mainly observed in cells that presented ASC specks. We found that punicalagin treatment did not alter PCNT loss induced by nigericin (Fig. 2I, J). We also found that in cells treated with ZVAD, and where ASC specks were able to form, no loss of PCNT occurred (Fig. 2I-K). Finally, treatment with MCC950 prevented assembly of the inflammasome indicated by the absence of ASC specks and no loss of PCNT stain was observed here (Fig. 2I, J).

We next tested if other NLRP3 inflammasome activators also triggered PCNT loss. First, we established that LPS priming of PMA-differentiated THP1 cells did not alter PCNT loss described in unprimed cells (Fig. S2). We found that LPS-primed THP1 cells still lost PCNT signal after 45 min nigericin treatment and this was prevented by the NLRP3 inhibitor MCC950 (Fig. S2). We then assessed the effect of LLOMe on PCNT loss. LLOMe mediates NLRP3-inflammasome activation by destabilizing the lysosomal membrane [32, 33]. We treated PMA-differentiated LPS-primed THP1 cells with LLOMe (1 mM, 1 h) and observed increased cell death and caspase-1 activity as expected. While ZVAD blocked caspase-1 activity it only partially blocked cell death (Fig. S3D, E). Like nigericin, LLOMe triggered PCNT loss, however, that was not prevented by treatment with ZVAD (Fig. S3A-C). Next, we tested the effect of NLRP3 activation via cell volume regulation [34] in PCNT regulation. For this we treated PMA-differentiated LPS-primed THP1 cells with a hypotonic solution for 1 and 3 h. We found that, as previously described, hypotonic shock led to caspase-1 activation and cell death at these time points (Fig. S3I, J). This was matched by loss of PCNT levels in the cell lysates at both time points (Fig. S3F-H).

In order to determine if PCNT loss occurred in a different cell type, we used murine BMDMs. As BMDMs require priming for appropriate NLRP3 inflammasome activation we treated these cells with LPS 1µg/ml for 4 h prior to nigericin treatment. We observed that as in THP1 cells, treatment of BMDMs with nigericin led to reduced PCNT expression and this was prevented by MCC950. However, unlike un-primed THP1 cells ZVAD failed to rescue the induced cell death and in these conditions PCNT levels were not recovered (Fig. S4). All of this suggests that PCNT loss is a general response when the NLRP3 inflammasome is activated.

### Nigericin leads to NLRP3 localization at centrosomal and non-centrosomal locations

As we had observed that nigericin triggered PCNT disruption in cells with an active inflammasome and given that the centrosome could be a place of assembly for the NLRP3 inflammasome, we next studied the relationship between the centrosome and NLRP3-location using a THP1 cell line stably expressing GFP-NLRP3 in an NLRP3 deficient background (THP1^GFP-NLRP3^) [35]. Upon NLRP3-activation with nigericin 30 min, to minimise centrosomal loss, we found that GFP-NLRP3 formed NLRP3-ASC-specks indicating active inflammasome platforms as expected, mainly at non-centrosomal locations. We also observed NLRP3 accumulation at the centrosome as GFP-NLRP3 co-localised with the pericentriolar material component PCNT, however these NLRP3-structures did not co-localise with ASC, suggesting this NLRP3 is non active. Both types of NLRP3-structures could be found simultaneously within the same cell (Fig. 3A). Nigericin treatment of THP1 cells expressing GFP alone in a WT background (GFP-THP1) did not lead to GFP re-location to the centrosome demonstrating that the observed effect is very likely driven by NLRP3 (Fig. S5). We next treated THP1^GFP-NLRP3^ cells with nigericin in the presence of the NLRP3 inhibitor MCC950 and ZVAD. We found that MCC950 prevented assembly of non-centrosomal NLRP3-ASC specks however did not alter the ability of NLRP3 to move to the centrosome (Fig. 3A). Pan-caspase inhibitor ZVAD did not prevent assembly of NLRP3-ASC-specks or NLRP3-association to the centrosome (Fig. 3A). In this experimental system we also detected PCNT loss after inflammasome activation as shown by quantification (Fig. 3B-D). Quantification of a time course of nigericin treatment in this cellular system showed similar results to those obtained in wild type THP1 cells indicating an increase in PCNT loss proportional to NLRP3-ASC-speck formation that was prevented by MCC950 (Fig. 3E). In these conditions, accumulation of NLRP3 at the centrosome occurred in the presence and absence of MCC950 (Fig. 3E, F). These data confirm that NLRP3 can be directed to the centrosome although in apparently a non-functional state, and that this directed localization occurs in parallel to inflammasome activation.

**Fig. 3.**
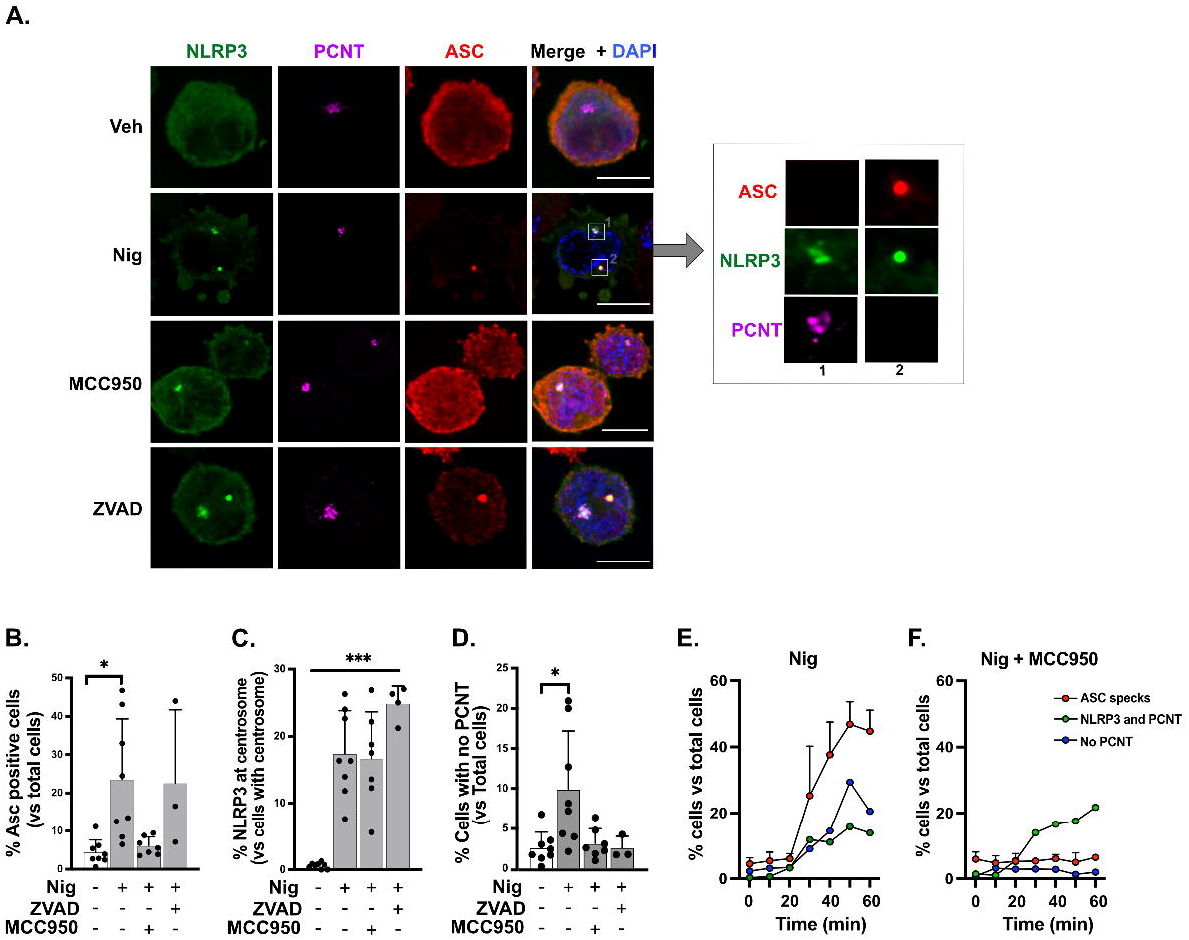
Nigericin leads to NLRP3 localisation at centrosomal and non-centrosomal locations. THP1^GFP-NLRP3^ cells were left untreated or treated with ZVAD (50 µM, 40 min) or MCC950 (10 µM, 15 min) prior to treatment with nigericin (10 µM, 45 min) to activate the NLRP3 inflammasome (A-D). Images show NLRP3 (green), ASC (Red) and PCNT (purple). Nuclei are shown in blue. ASC speck positive cells were quantified and plotted as percentages versus total number of cells (B). Percentages of cells with NLRP3 at centrosome relative to cells with centrosome and cells with no PCNT in total cells were calculated respectively (C, D). THP1^GFP-NLRP3^ cells were left untreated or treated with MCC950 (10 µM, 15 min) before in response to nigericin (10 µM) at different time points as indicated (E, F). Percentages of cells with ASC specks, or with NLRP3 and PCNT, or no PCNT in total cells were calculated. 300 cells were counted and analyzed per experiment, N=3, Independent experiments.

### PCNT loss induced by NLRP3 inflammasome is dependent on caspase-1 activation

Having shown that PCNT loss depends upon NLRP3 activation and that it can be prevented by pan caspase inhibitor ZVAD in THP1 cells we next considered the specific effect of caspase-1 in PCNT loss, initially using the caspase-1 inhibitor YVAD (36). YVAD pre-treatment blocked caspase-1 activity and IL-18 release but only partially decreased cell death (Fig. 4D-F) induced by nigericin. This is reflected in YVAD treatment partially rescuing the downregulation of PCNT induced by nigericin in vehicle treated cells (Fig. 4A-C). To further confirm the contribution of caspase-1 to PCNT degradation we used THP1 cells deficient for caspase-1 (THP1^Caspase1-/-^ cells). We found that nigericin treatment of these cells did not induce pyroptosis or IL-18 release (Fig.4H-J). When looking at PCNT expression we found that in the absence of caspase-1 PCNT levels were not reduced in response to nigericin. However, we detected a cleaved band of around 200kDa that would correspond to the size of caspase-3 mediated cleavage of PCNT previously described [36] since nigericin triggered activation of caspase-3 in these cells (Fig. 4G). Our data suggest that caspase-1 activation contributes to PCNT degradation when NLRP3 inflammasome is activated by nigericin in THP1 cells.

**Fig. 4.**
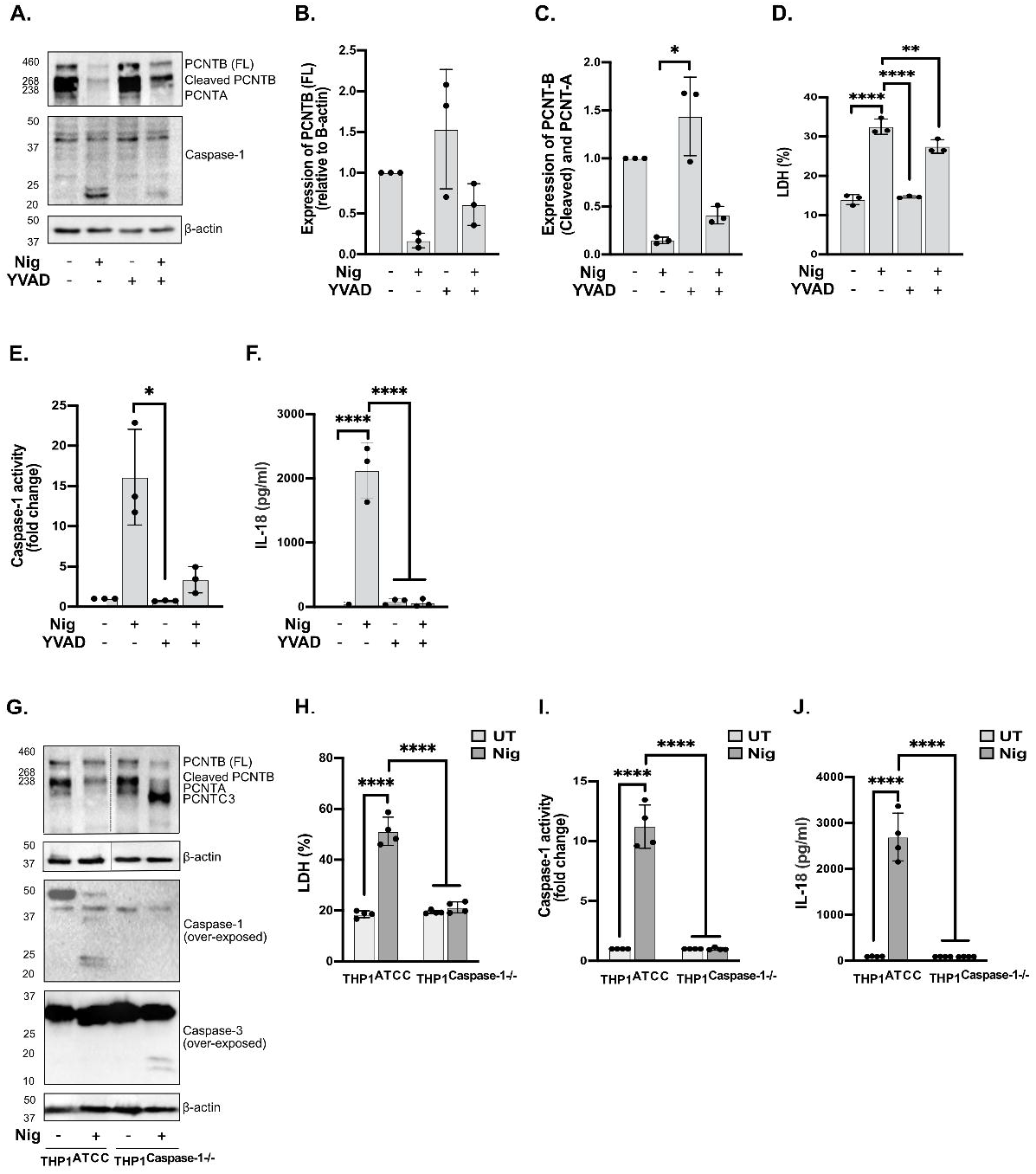
PCNT loss induced by NLRP3 inflammasome is dependent on caspase-1 activation. THP1^Null2^ cells were left untreated or treated with YVAD (50µM, 40 min), then stimulated with nigericin (10 µM, 45 min) (A-F, N=3). Lysates were analyzed for PCNT as well as loading control β-actin by western blot (A). Relative expression of full length PCNT-B and cleaved PCNT-B/PCNT-A compared to the β-actin was quantified respectively (B, C). Cell death (D), caspase-1 activity (E) and IL-18 (F) were measured as described above. THP1^ATCC^ and THP1^Caspase-1-/-^ cells were directly stimulated with nigericin (10 µM, 45 min) (G-J, N=4). Lysates were analyzed for PCNT, caspase-1 and caspase-3 as well as loading control β-actin (G). Cell death (H), caspase-1 activity (I) and IL-18 (J) were measured as described above. Independent experiments. For multiple comparisons, one-way ANOVA with the Dunnett’s test for YVAD in THP1^Null2^ cells and two way ANOVA with the Tukey’s test for comparing nigericin treated THP1^ATCC^ and THP1^Caspase-1-/-^ cells were applied. Data was shown as mean ± S.D., *p < 0.05, **p < 0.01, ***p < 0.001, ****p < 0.001 were considered statistically significant.

### Proteasomal and lysosomal activity does not affect NLRP3-induced PCNT loss

We next wanted to understand what controlled the described loss of PCNT. As proteasome inhibition regulates levels of PCM proteins including PCNT [36] we first tested the role of the proteasome in PCNT loss. For this we treated THP1 PMA differentiated cells with proteasome inhibitor MG132 (10 µM) for 2 h prior to nigericin treatment. We found that nigericin treatment increased proteasome activity, and that this was blocked by MG132 treatment (Fig. S6A). We found that total PCNT levels were increased when cells were treated with MG132 alone, indicating PCNT regulation by the proteasome (Fig. 5A). However, when treated with nigericin, these cells still lost PCNT in the presence of MG132 (Fig. 5A). Overall, these data suggest that PCNT degradation induced by nigericin is not mediated by proteasomal regulation.

**Fig. 5.**
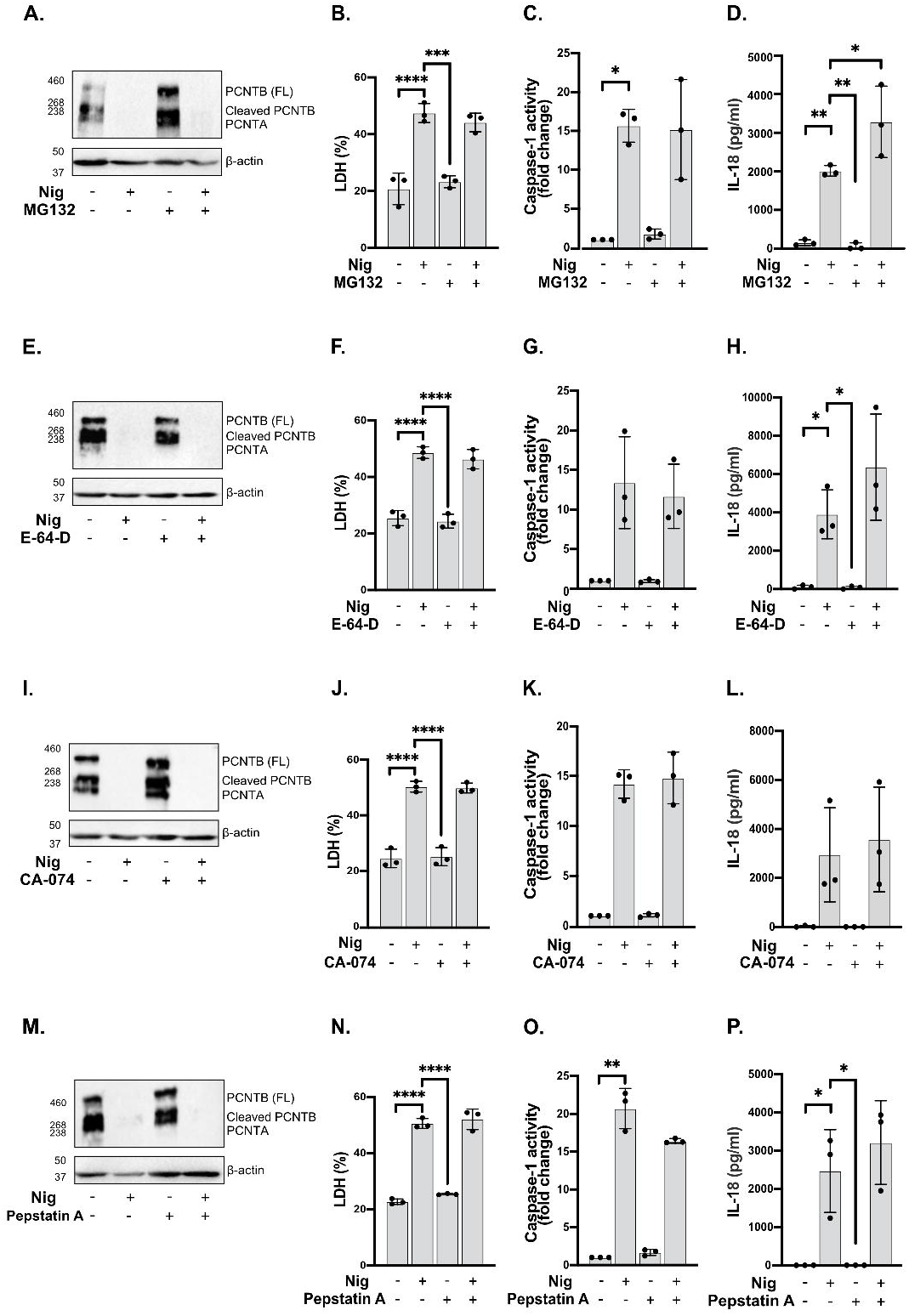
Inhibition of proteasomal and lysosomal activity does not affect NLRP3-induced PCNT loss. PMA differentiated THP1^ATCC^ cells were left untreated or treated with MG132 (10 µM, 2 h), or E-64-D (20 µM, 2 h), or Ca-074Me (50 µM, 15 min), or pepstatin A (10 µM, 15 min) before stimulation with nigericin (10 µM, 45 min) (A-P). Lysates were analyzed for PCNT as well as loading control β-actin by western blot (A, E, I, M). Cell death (B, F, J, N), caspase-1 activity (C, G, K, O) and IL-18 (D, H, L, P) were measured as described above. Data was shown as mean ± S.D., *p < 0.05, **p < 0.01, ***p < 0.001, ****p < 0.001 were considered statistically significant.

We next explored the links between lysosome and PCNT loss. Lysosomal cathepsins have been linked to inflammasome activation mediated by nigericin [32]. We tested the role of cathepsins by using the cathepsin inhibitor E-64-D (20 µM) for 2 h, the cathepsin B inhibitor CA-074 (50 µM) for 15 min, or the cathepsin D inhibitor pepstatin A (10 µM) for 15 min before nigericin stimulation. We confirmed activity of these inhibitors as we found that E-64-D and CA-074 decreased cathepsin B activity, and pepstatin A reduced cathepsin D activity in THP1 cells (Fig. S6B-D). We found that these inhibitors did not affect cell death, caspase1 activity or IL-18 release levels induced by nigericin (Fig. 5F-H, J-L, N-P). Similar to what was observed with MG132 treatment, nigericin stimulation resulted in PCNT degradation even in the presence of these inhibitors (Fig. 5E, I, M) suggesting that cathepsins are not required for the PCNT degradation.

### GSDMD is required for PCNT disruption triggered by pyroptosis but not by apoptosis

Caspase-1 activation triggered by NLRP3 inflammasome results in GSDMD cleavage and consequently assembly of GSDMD pores in the plasma membrane triggering pyroptosis [37]. These pores have also been described as conduits for release of IL-1β. To examine the link between GSDMD and pyroptosis in PCNT loss we compared expression of PCNT in THP1^WT^ and THP1^GSDMD-/-^ cells pre-treated with vehicle or ZVAD and followed by nigericin activation (10 µM, 45 min). GSDMD deficiency led to reduced cell death (Fig. 6A) and extracellular caspase-1 activity (Fig. 6B) compared to WT cells, as caspase-1 release was prevented in these cells. Despite the presence of active caspase-1, nigericin treatment of THP1^GSDMD-/-^ did not lead to PCNT loss as in WT cells, but to PCNT processing indicative of caspase-3 mediated cleavage (Fig. 6C) (as in caspase-1 deficient THP1 cells) given that GSDMD deficiency leads to a switch from pyroptosis to apoptosis in response to nigericin with activation of caspase-3 [23, 24]. Activation of caspase-3 and caspase-1 was confirmed in the lysates of WT and GSDMD deficient cells (Fig. 6D). We confirmed that the observed caspase-3 mediated PCNT cleavage was prevented by ZVAD (Fig. 6C) and that was also blocked by caspase-3 inhibitor Z-DVED confirming this processing is caspase-3 dependent (Fig. 6E-G). Furthermore, nigericin treatment in the presence of Z-DVED not only prevented caspase-3 mediated PCNT cleavage, but partially recovered the PCNT loss as observed in WT cells.

**Fig. 6.**
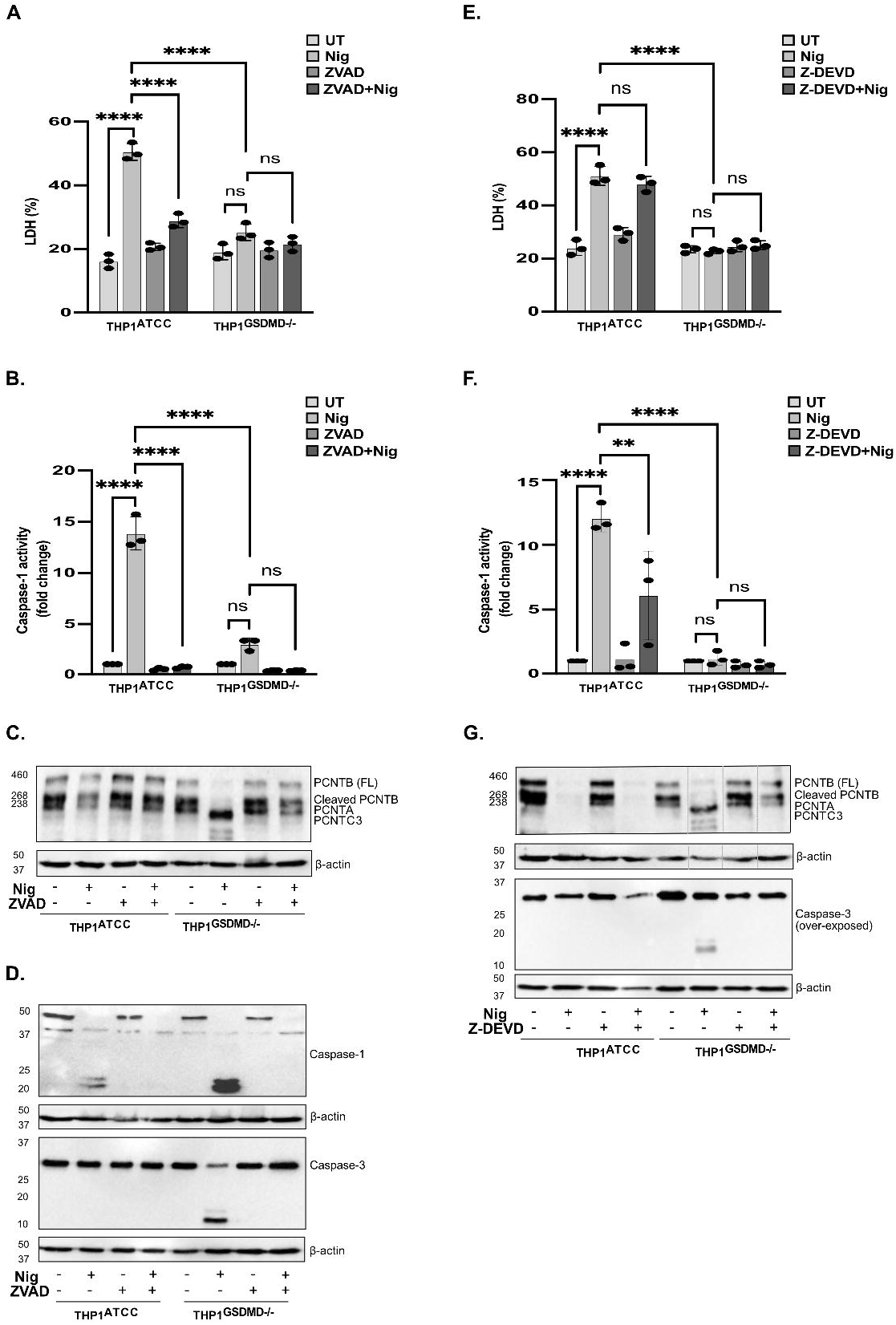
GSDMD is required for PCNT disruption triggered by pyroptosis but not by apoptosis. THP1^ATCC^ and THP1^GSDMD-/-^ cells were left untreated or treated with ZVAD (50 µM, 40 min) or Z-DEVD (20 µM, 2 h), after which cells were stimulated with nigericin (10 µM, 45 min) to activate the NLRP3 inflammasome (A-G). Caspase-1 activity was measured by caspase-1 assay and shown as fold change relative to control (A, E). Cell death was measured as described above and shown as percentage relative to total cell death (B, G). Lysates were analyzed for PCNT, caspase-1 and caspase-3 as well as loading control β-actin (C, D, G). Data was shown as mean ± S.D., *p < 0.05, **p < 0.01, ***p < 0.001, ****p < 0.001 were considered statistically significant.

## Discussion

In recent years the centrosome has been proposed as an important player in NLRP3 inflammasome activation by acting as a cellular location for inflammasome assembly [6, 7] as well as for regulating activation via centrosomal proteins like PLK1 [9] and PLK4 [10]. Our data adds to this knowledge by showing that the centrosome is disrupted after inflammasome activation. Here we have found that NLRP3 inflammasome activation by different triggers, nigericin, LLOMe and hypotonicity, leads to loss of centrosomal proteins and centrosomal disorganization. This disruption is dependent on caspase-1 and GSDMD and we show that pyroptosis, but not membrane rupture, is driving this centrosomal loss in human and murine macrophages.

Pyroptosis is a characteristic type of cell death driven by inflammasome activation [22]. GSDMD and ninjurin-1 are two important mediators in this cellular mode of cell death. GSDMD cleavage and pore formation in response to inflammasome activation are required for IL-1β and IL-18 release [38, 39], as well as driving pyroptotic cell death, while ninjurin-1 is responsible for the plasma membrane rupture that follows GSDMD pore formation [40, 41]. Our observation that treatment with the membrane stabiliser punicalagin did not prevent centrosomal loss or inflammasome activation, but just cytokine release, suggests that this PCNT loss is intrinsically associated to the pyroptotic process and not membrane rupture. GSDMD deficient cells treated with nigericin, are still able to form an active inflammasome, like cells treated with punicalagin. However, GSDMD deficient cells do not die of pyroptosis but apoptosis mediated by caspase-3 despite caspase-1 still being active. In these conditions, PCNT was prevalently cleaved by caspase-3 suggesting that although caspase-1 is involved in centrosomal disruption, this is not directly mediating PCNT loss.

We observed that, except in the case of punicalagin or GSDMD deficient cells, PCNT-loss was proportional to the release of LDH that occurred after activation. This was more obvious after the use of ZVAD or YVAD. We found that in LPS-treated cells (THP1 and BMDMs) ZVAD was active and able to block IL-18 and IL-1β release, however, had no major effect on LDH release. This differential effect of YVAD and ZVAD has been previously described [42]. Here the authors showed that these compounds fail to block the cleavage of the effector molecule GSDMD by caspase-1 and hence prevent LDH release [42] while still able to block cleavage of IL-1β and IL-18. This provides yet another link between the pyroptotic process and PCNT loss. Why we mainly observed this in LPS treated cells we still do not understand and would require further studies.

We have found that nigericin treatment leads to the accumulation of non-active GFP-NLRP3 at the centrosome and active NLRP3-GFP at non-centrosomal locations. These observations agree with a recent report showing that NLRP3 tagged with neon Green (NLRP3-mNG) aggregate at both centrosomal and non-centrosomal localizations in THP1 cells [43]. The centrosome acts as a signalling hub where proteins come in and out to tightly regulate cellular processes such as DNA damage [15] or cell cycle entry [16]. Hence one could think that NLRP3 accumulates at the centrosome before becoming fully functional as a way of controlling unwanted excessive activation. Whether NLRP3 is then degraded or released to form an active inflammasome is currently unknown. When we looked at endogenous ASC-speck formation we found that the centrosome was not the predominant ASC-speck localization, although could be observed in some cells. Although this differs from the reports of Li [6] and Magupalli [7], this is consistent with our data using GFP-NLRP3 as well as recent work from Liu Y, et al. (2023), which found that although ASC-specks are closer to the microtubule organizing centre (MTOC), they do not co-localize with this organelle markers [44]. It is possible that localization of ASC-specks at the centrosome is very transient and considering the disruption of the centrosome described here it might be difficult to detect such cellular positioning.

Centrosome plasticity is manifested during inflammation. LPS priming of macrophages induces recruitment of PCM components such as PCNT and γ-tubulin to the centrosome and is important for cytokine secretion [19]. This process reminds of centrosome amplification but occurs during interphase and independently of PLK1 [19]. Similarly in microglia LPS leads to the recruitment of microtubule nucleating material to the centrosome [45]. This might be linked to the ability of LPS to arrest cells in G0/1 state, altering cell cycle [18, 46] and hence centrosomal composition [19]. Centrosomal restructuring has also been reported in monocytes from febrile patients. This occurred by proteasomal mediated degradation at the centrosome induced by heat shock. Interestingly, this process mediated by Hsp70, was reversible [17]. Although authors propose that this is a process important for the immune response, whether macrophages function during inflammation is affected when centrosome is re-structured remains to be assessed. Heat shock induced centrosomal disruption resembles what we have observed in macrophages after inflammasome activation. Our study adds to the evidence that alterations on centrosomal composition occur during the inflammatory response. In our experimental conditions we observed an increase in proteasome activity after nigericin treatment that was prevented by proteasome inhibitor MG132. However, this did not prevent centrosomal disruption indicating that the mechanisms of regulation between these two processes are different. Proteasomal proteins can traffic in and out of the centrosome and can be degraded via other pathways such as autophagy and lysosomal degradation [47]. However, we failed to impair centrosomal loss or inflammasome activation by blocking either proteasome or lysosomal cathepsins. This suggests that there must be an alternative mechanism of degradation or compensatory pathways.

Microtubules are important for inflammasome function. Trafficking along microtubules to different locations in the cells, including the centrosome, is important for inflammasome activation [6, 7]. Microtubule remodelling is also an important event in cell death. During apoptosis, microtubules are reformed organizing an apoptotic microtubule network important for maintaining plasma membrane integrity and cell morphology during the execution phase of apoptosis. Disruption of this network is however linked to secondary necrosis [48]. Disorganization of the cytoskeleton also occur during pyroptosis. Infection with Salmonella typhimurium induced loss of cytoskeletal marker α-tubulin, which was prevented in caspase-1 knock out cells [49]. Calcium entry triggered by inflammasome activation leads to calpain-mediated vimentin cleavage and release and disruption of intermediate filaments contributing to loss of cytoskeleton stability in THP-1 cells [38]. However, neither calpain inhibition nor EGTA treatment to chelate calcium prevented loss of actin filaments, microtubules, or nuclear lamina indicating that these proteins are regulated by a different mechanism that occurs in parallel to vimentin cleavage. Our data add to this showing that not only microtubules, but also the main microtubule organising centre is disrupted upon pyroptosis, suggesting that maybe centrosome disruption is the first step leading to cytoskeletal disassembly and rupture. This centrosomal disorganization could facilitate release of ASC-speck to the extracellular environment to propagate inflammation to neighbouring cells [50, 51].

Despite the importance of the centrosome in cell homeostasis and recent advances around inflammasome activation and the centrosome, little is known about the role of this organelle in macrophage function during inflammation. Here we have reported the disruption of the centrosome upon NLRP3-inflammasome activation. However, when does this centrosome remodelling commences and how this is regulated, we still do not understand. Additionally, how LPS, or other priming signals, alter centrosomal composition and how this contributes to different priming events, have not yet been explored. Moreover, the implications for NLRP3 presence at the centrosome after sensing NLRP3-activators are not understood and this deserves further studies.

## Contributions

Performed experiments and sample collection: S.B. and F.M.S.

Experimental design: D.B. and GLC

Supervision: D.B. GLC

Writing of manuscript: SB, FMS, DB and GLC.

## Acknowledgments

This work was supported by a Presidents Doctoral Scholarship (The University of Manchester) to Siyi Bai, a Wellcome Trust and Royal Society Henry Dale Fellowship to G. L-C. (104192/Z/14/Z), a Medical Research Council grant (MR/T016043/1) awarded to G.L.C. and by a Medical Council Research grant awarded to DB (MR/T016515/1). The Bioimaging Facility microscopes used in this study were purchased with grants from BBSRC, Wellcome, and the University of Manchester Strategic Fund.

## Conflict of Interest

The authors declare that they have no conflicts of interest regarding this study.

## Notes

### Competing Interest Statement

The authors have declared no competing interest.

