## Supplemental figures for "Pyroptosis leads to loss of centrosomal integrity in macrophages"

Fig S1

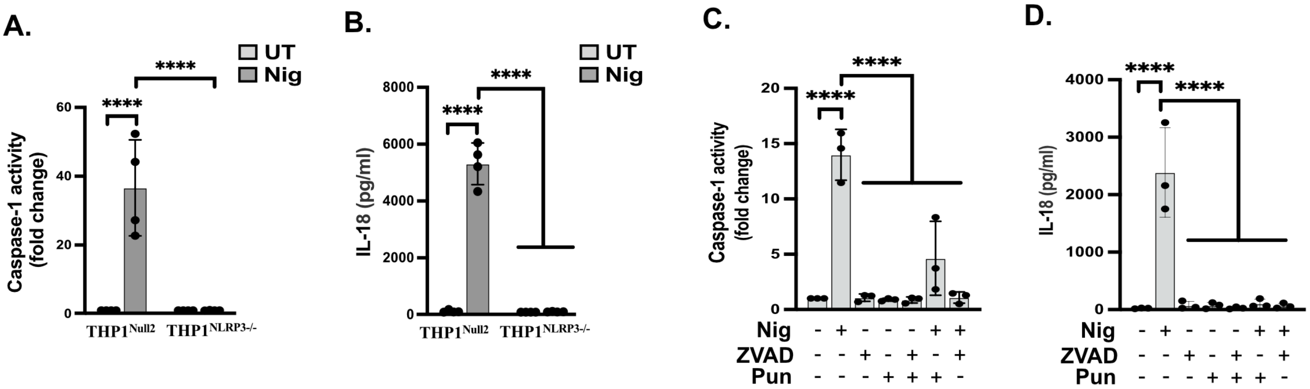


**Fig.S1 Caspase-1 activity and IL-18 release in THP1 with impaired NLRP3 activation-Related to figure 1.**

THP1$^{Null2}$ and THP1$^{NLRP3-/-}$ cells were stimulated with nigericin (10 µM, 45 min) (A-B). THP1$^{Null2}$ cells were left untreated or treated with punicalagin (50 µM, 15 min) or ZVAD (50 µM, 40 min), after which cells were stimulated with nigericin (10 µM, 45 min) to activate the NLRP3 inflammasome (C-D). Caspase-1 activity was measured by caspase-1 assay and shown as fold change relative to control (A, C). IL-18 was measure by ELISA (B, D).

Fig S2

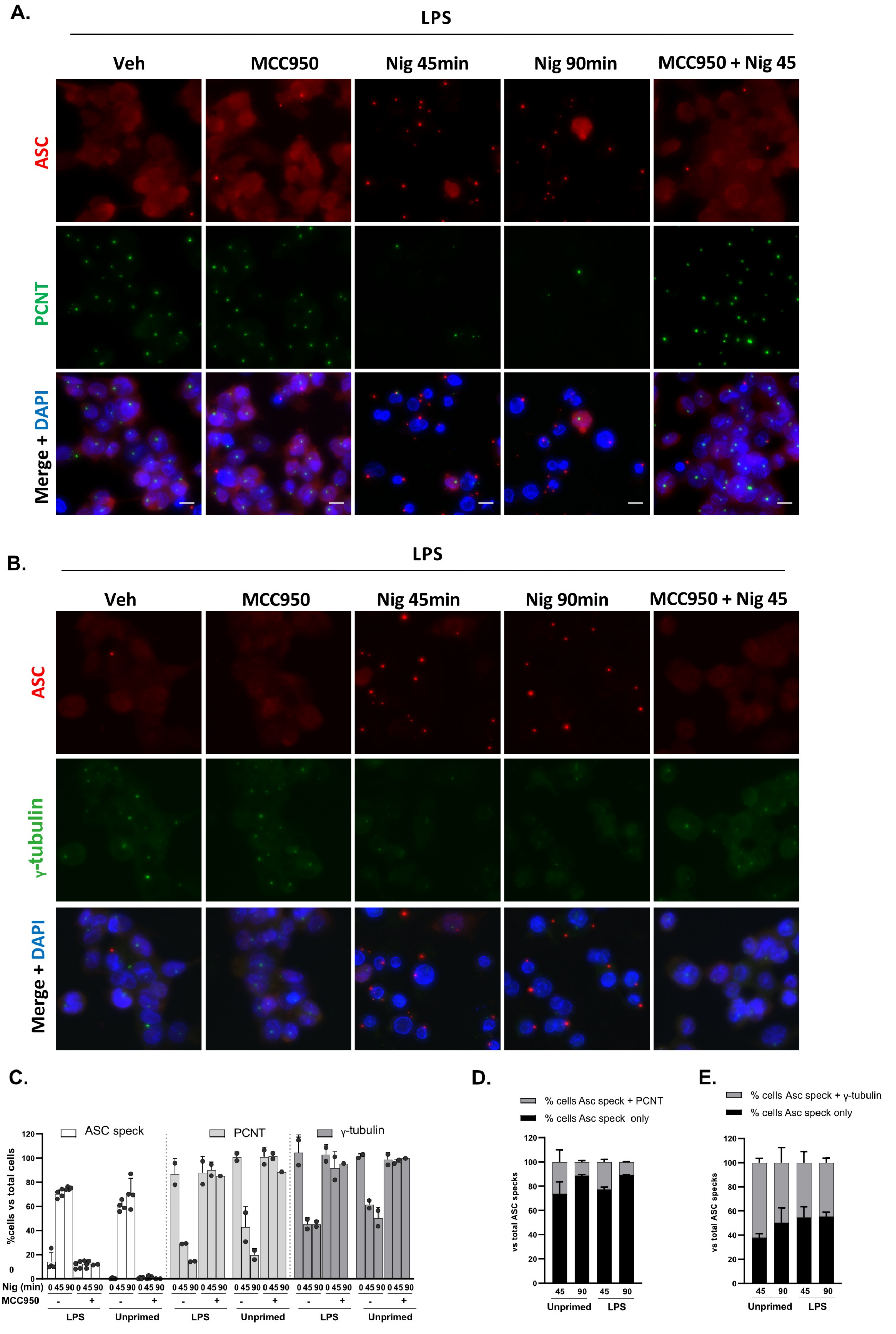

A.
LPS
Veh
MCC950
Nig 45min
Nig 90min
MCC950 + Nig 45
ASC
PCNT
Merge + DAPI
B.
LPS
Veh
MCC950
Nig 45min
Nig 90min
MCC950 + Nig 45
ASC
γ-tubulin
Merge + DAPI
C.
ASC speck
PCNT
γ-tubulin
%cells vs total cells
Nig (min)
MCC950
LPS
Unprimed
D.
% cells Asc speck + PCNT
% cells Asc speck only
vs total ASC specks
Unprimed
LPS
E.
% cells Asc speck + γ-tubulin
% cells Asc speck only
vs total ASC specks
Unprimed
LPS

**Fig.S2 Priming does not alter PCNT loss caused by nigericin in THP1 cells.**

THP1 [ATCC] cells were primed with LPS (1µg/ml, 4 h) or not, then treated with MCC950 (10 µM, 15 min), after which cells were stimulated with nigericin (10 µM) for 45 or 90 min (A-D). Immunofluorescence was used to analyze PCNT, $\gamma$-tubulin and ASC (A). Percentages of ASC speck, PCNT or $\gamma$-tubulin positive cells relative to total cells were calculated (B). Percentages of both ASC speck and PCNT or only ASC speck positive cells in total ASC positive cells (C) and of both ASC speck and $\gamma$-tubulin or only ASC speck positive cells in total ASC positive cells (D) were calculated. 300 cells were analyzed per experiment. Independent experiments, n=3.

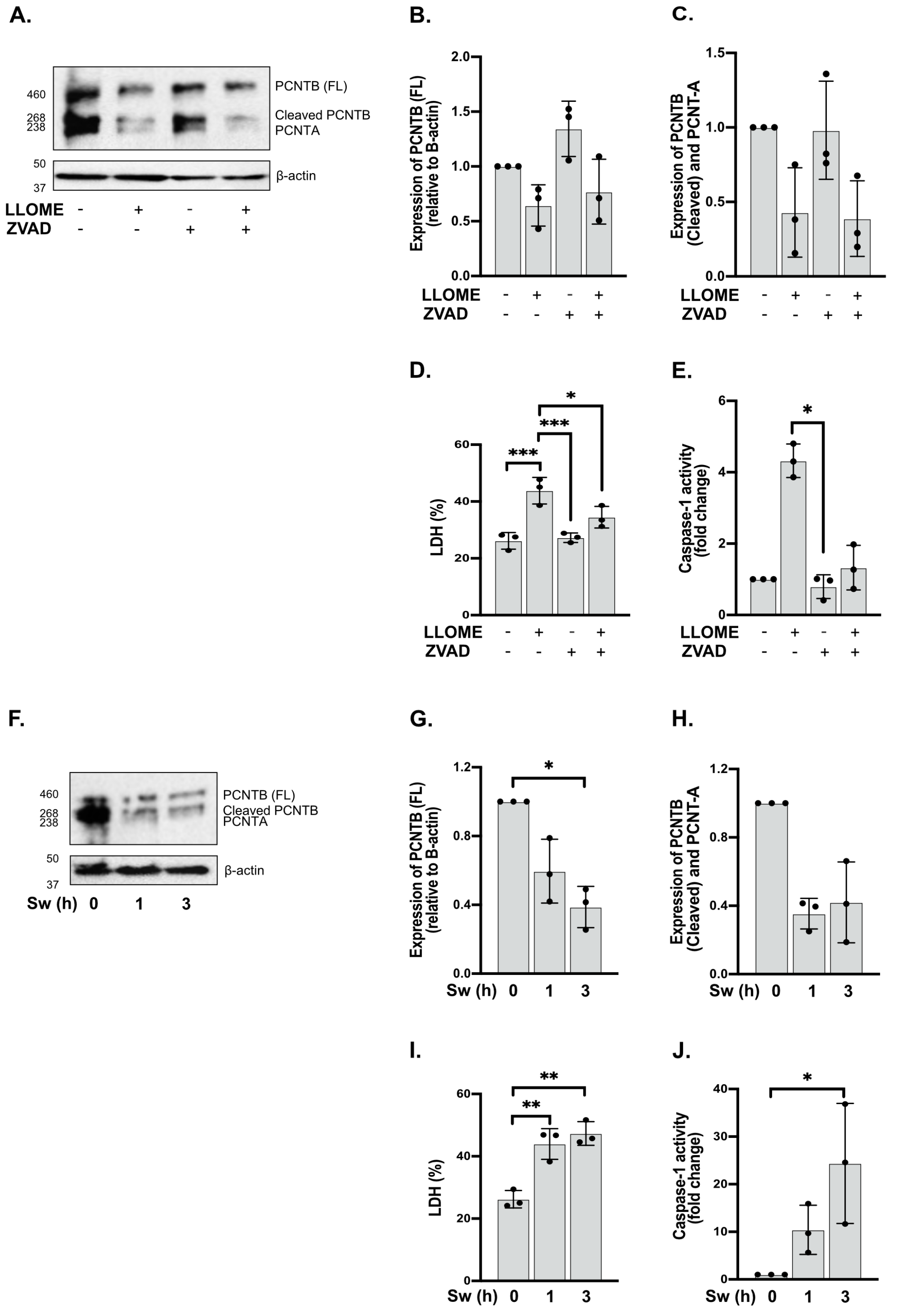


**Fig.S3 LLOMe and cell volume regulation trigger PCNT degradation.**
THP1[ATCC] cells were left untreated or treated with ZVAD (50 µM, 40 min), then treated with LLOMe (1 mM, 50 min) to activate the NLRP3 inflammasome (A-E, N=3). Lysates were analyzed for PCNT as well as loading control β-actin (A). Relative expression of full length PCNT-B and cleaved PCNT-B/PCNT-A compared to the β-actin was quantified respectively (B, C). Cell death was measured by LDH assay and shown as percentage relative to total cell death (C). Caspase-1 activity was measured by caspase-1 assay and shown as fold change relative to control (D). THP1[ATCC] cells were left untreated or treated with a hypotonic buffer for 1 or 3 h to activate the NLRP3 inflammasome (F-J, N=3). Lysates were analyzed for PCNT as well as loading control β-actin (F). Relative expression of PCNT (G-H), cell death (I) and caspase-1 activity (J) were quantified or measured as described above.

Fig S4

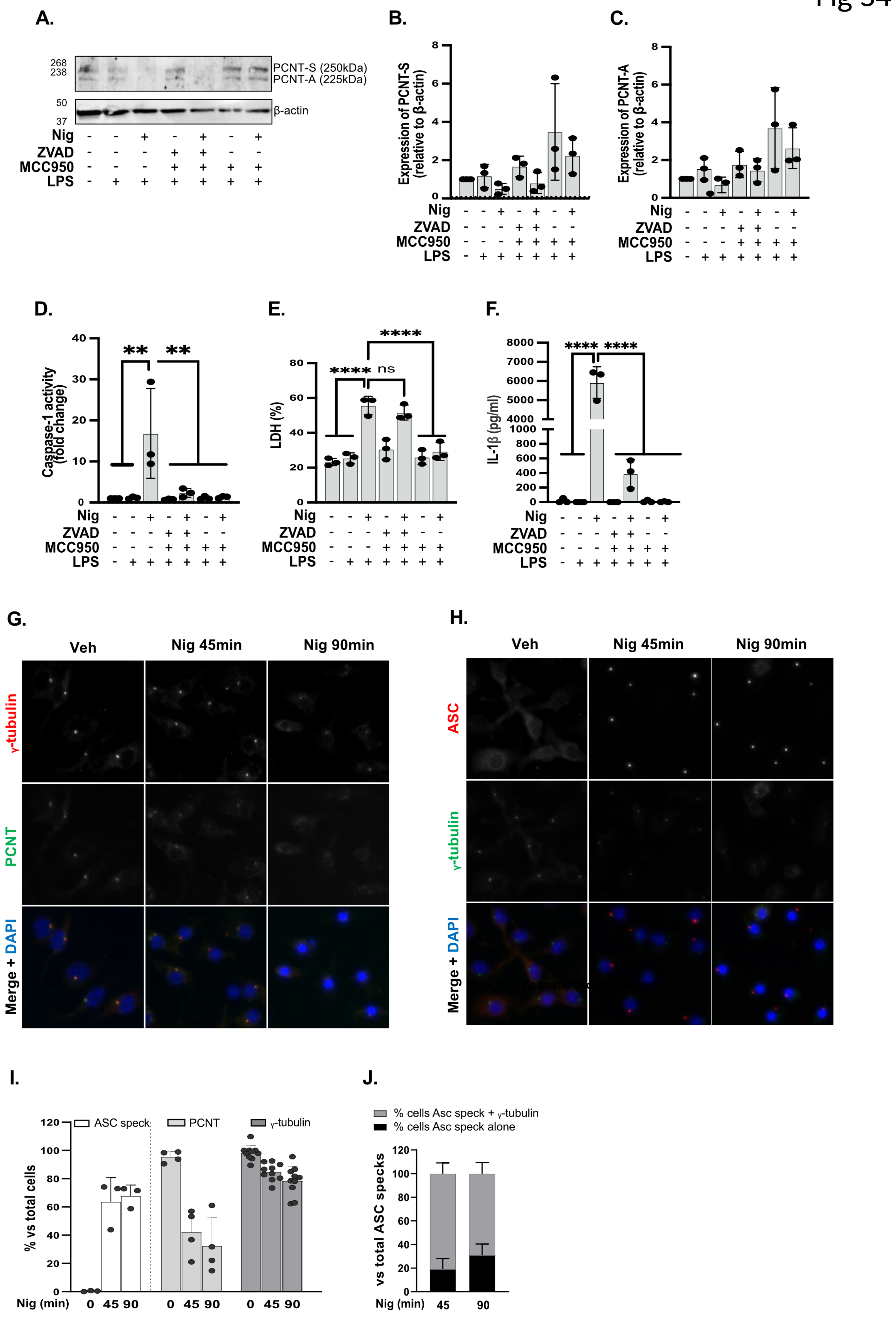

A.
268
238
PCNT-S (250kDa)
PCNT-A (225kDa)
50
37
β-actin
Nig
ZVAD
MCC950
LPS
B.
Expression of PCNT-S (relative to β-actin)
C.
Expression of PCNT-A (relative to β-actin)
D.
Caspase-1 activity (fold change)
E.
LDH (%)
ns
F.
IL-1β (pg/ml)
G.
Veh
Nig 45min
Nig 90min
γ-tubulin
PCNT
Merge + DAPI
H.
ASC
γ-tubulin
Merge + DAPI
I.
ASC speck
PCNT
γ-tubulin
% vs total cells
Nig (min)
J.
% cells Asc speck + γ-tubulin
% cells Asc speck alone
vs total ASC specks
Nig (min)

**Fig.S4 Nigericin triggers PCNT loss in BMDMs.**
LPS (1μg/ml, 4 h) primed BMDMs were left untreated or treated with MCC950 (10 μM, 15 min) or ZVAD (50 μM, 40 min), then cells were stimulated with nigericin (10 μM, 45 min) (A-F, N=3). Lysates were analyzed for PCNT as well as loading control β-actin by western blot (A). Relative expression of full length PCNT-S (250 kDa) and PCNT-A (225 kDa) compared to the β-actin was quantified respectively (B, C). Cell death (D), caspase-1 activity (E) and IL-18 (F) were measured. LPS (1μg/ml, 4 h) primed BMDMs were left untreated or treated with nigericin (10 μM) for 45 or 90 min to activate the NLRP3 inflammasome (G-J). Immunofluorescence was used to analyze PCNT, γ-tubulin and ASC (G, H). Percentages of ASC speck, PCNT or γ-tubulin positive cells relative to total cells (I) and both ASC speck and γ-tubulin or only ASC speck positive cells in total ASC positive cells (J) were calculated. 300 cells were counted and analyzed per experiment. Independent experiments, n=3.

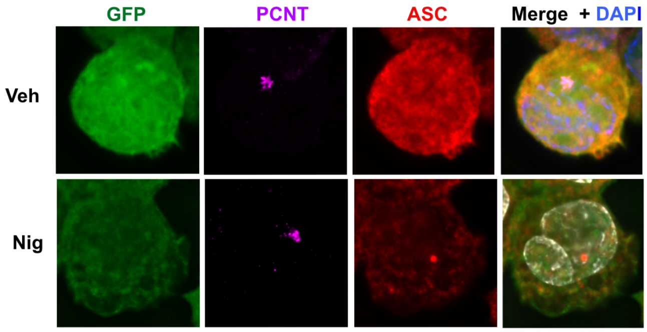


**Fig.S5 GFP alone does not re-locate to the centrosome with nigericin stimulation in THP1 cells.**
GFP-THP1 cells were left untreated or treated with nigericin (10 μM, 45 min) (A, n=3). Immunofluorescence was used to analyze GFP, PCNT and ASC (A).

Fig S6

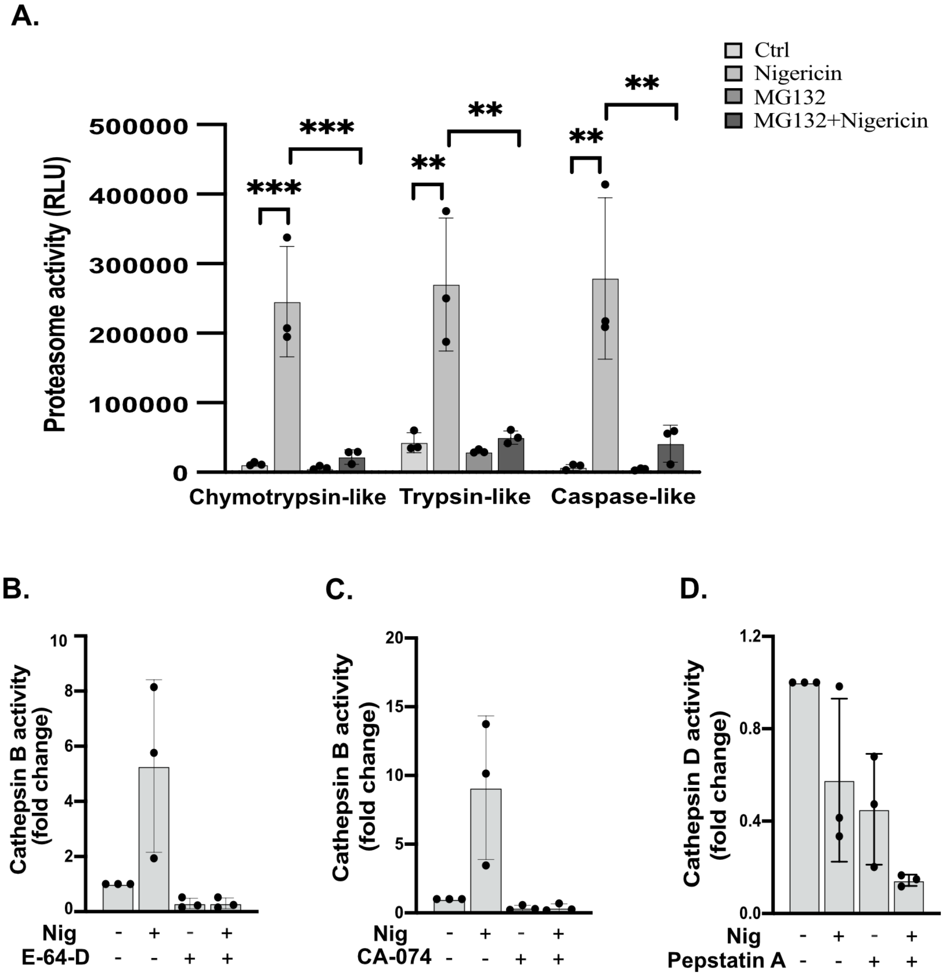


**Fig.S6 Proteasome, CathepsinB and Cathepsin D activities during the NLRP3 activation in THP1 cells-Related to figure 5.**

PMA differentiated THP1$^{ATCC}$ cells were left untreated or treated with MG132 (10 µM, 2 h), or E-64-D (20 µM, 2 h), or Ca-074Me (50 µM, 15 min), or pepstatin A (10 µM, 15 min) before stimulation with nigericin (10 µM, 45 min) to activate the inflammasome (A-D). Proteasome activity was measured, and luminescence was recorded as relative light units (RLU) (A). The activities of cathepsin B and cathepsin D were measured and shown as fold change from the untreated cells control for all experimental groups (B-D).
